# Evidence from metagenomic study indicate that subclinical mastitis may have a different pathological origin than clinical mastitis

**DOI:** 10.1101/2024.05.24.595548

**Authors:** Subhrajit Bhar, Tungadri Bose

## Abstract

Bovine mastitis is one of the main causes of low milk production, resulting in significant economic losses for the dairy industry. Therefore, the industry will benefit from the development of strategies for the timely diagnosis of bovine mastitis, especially the sub-clinical sub-type. Here, we analysed the milk metagenome of healthy cows and cows suffering from various forms of mastitis, viz. clinical, sub-clinical and chronic/ recurrent sub-types. We identified *Neisseria*, *Eubacterium* and *Streptococcus* as the key drivers of the change in microbial community structure from a healthy state to a clinical mastitis state. Our results also indicate that the microbiota composition and the probable cause of clinical and recurrent bovine mastitis may not be the same as that of sub-clinical mastitis. Further, the sensory protein load in the sub-clinical mastitis sub-group differed significantly from the other studied categories, wherein *Achromobacter*, *Dickeya*, *Pectobacterium* and *Raoultella* were identified as the discriminatory features. We also propose ML-based classifiers to screen for bovine mastitis using milk metagenomic samples. The principles elucidated here through the study of mastitis in cows can be applied to other animals and hopefully will benefit the entire dairy industry.

## Introduction

Bovine mastitis (BM) is one of the leading causes of decrease in milk yield resulting in significant economic losses to the dairy industry (Azooz et al., 2020; Nikkhah et al., 2021). BM may be caused by physical trauma or microbial infection of the mammary gland, resulting in inflammation of udder tissue (Cheng and Han, 2020). Based on the degree of inflammation and/or frequency of re-occurrence, BM can be classified into clinical (CM), sub-clinical (SCM) and chronic/ recurrent (RCM) sub-types. While CM and RCM are symptomatic, SCM shows no evident abnormalities but results in significant decrease in milk production (Cheng and Han, 2020; Romero et al., 2018). While common bacterial pathogens like *Escherichia coli*, *Staphylococcus aureus* and *Streptococcus uberis* are known to cause BM, SCM can be caused by less common genera such as *Mycoplasma* and *Corynebacterium* as well (Cheng and Han, 2020; Cobirka et al., 2020). Timely diagnosis of SCM is therefore often challenging (Kour et al., 2023) and strategies to effectively manage SCM, in particular, would hugely benefit the dairy industry (Romero et al., 2018).

Bovine (cow) udder contains a natural community of microbes, and the collective composition of these microbes constitutes the bovine mammary gland microbiota, dysbiosis of which is likely to play a role in determining mastitis susceptibility (Derakhshani et al., 2018). Being an invasive process, (periodic) collection of the udder microbiota for analysis can be challenging. Notably, the microbiota of the bovine milk is highly similar to the bovine mammary gland (udder) microbiota (Taponen et al., 2019) and high-throughput next-generation sequencing techniques have enabled the study of the microbial communities in bovine milk (Addis et al., 2016; Hoque et al., 2020b, 2020a, 2020a; Taponen et al., 2019). Therefore, analysing the bovine milk microbiota may offer insights on the udder microbiota status and help in early assessment of the risk of developing BM. Earlier studies have already reported difference in the microbiota, in terms of microbial abundances and the functions encoded by them, in milk samples collected from healthy and mastitis affected cows (Hoque et al., 2020b, 2019). Studies have also shown dissimilarities in the composition of microbiota obtained from milk of cows affected with different sub-types of mastitis (viz., clinical, sub-clinical and recurrent/ chronic) (Bhatt et al., 2012; Hoque et al., 2019; Oikonomou et al., 2014)[14-16]. Whether these differences in microbiota composition can be used in designing a biomarker for early assessment of BM is worth exploring. In this context, it may be noted that the signal transduction system in bacteria is a key mechanism involved in acclimatization of the microbes to its ecological niche (Bose et al., 2017; Capra and Laub, 2012; Huang et al., 2015). Presence of the appropriate repertoire of sensory proteins is expected to benefit the microbes in adapting to the changing microenvironment of their host, and therefore their abundance in the metagenome may be associated with the onset and progression of a disease (Bhar et al., 2022). It would therefore be interesting to explore if any differences in the Sensory Protein Load (SPL) exists between the microbiota samples collected from milk of healthy versus diseased cows. Moreover, analysis of community interaction patterns between the microbes comprising the milk microbiota are largely missing. Therefore, it would also be interesting to inspect whether the community interaction pattern varies between the milk samples collected from healthy cows and those affected with mastitis.

To obtain perspectives into the above-mentioned unexplored areas, publicly available data containing milk microbiota samples from healthy cows (H), and cows affected by clinical mastitis (CM), sub-clinical mastitis (SCM), and recurrent/ chronic mastitis (RCM) were obtained (Hoque et al., 2020b, 2020a, 2019) for analysis. Distinct differences in the microbial community interaction patterns were observed between the H and CM samples. The changes in the community interaction in the diseased (CM) samples were driven by *Neisseria*, *Eubacterium* and *Streptococcus*. In addition, the SCM samples were found to be depleted in the abundance of known BM causing pathogens like *Acinetobacter* and *Salmonella* when compared to CM and RCM samples. Further, in terms of SPL, significant differences were noted in SCM samples when compared to the other two disease categories (CM and RCM) as well as the healthy (H) sub-group. ML-models with the capability to distinguish between milk metagenomic samples from health cows and those suffering from BM (viz., CM, SCM and RCM) is also presented. Overall, the data presented is expected to help in designing of tools for timely management of BM, thus benefitting the dairy industry.

## Results

### Clustering of the metagenomic samples were based on disease status

A dataset comprising of metagenomic sequences from 42 bovine milk samples collected across five different geographical locations (i.e., Chittagong, Dhaka, Gazipur, Manikgonj and Sirajgonj) and from cows belonging to four different health status (viz., healthy – H; clinical mastitis – CM; sub-clinical mastitis – SCM; and recurrent or chronic mastitis – RCM) was used for analysis. Supplementary Table 1 shows the distribution of the samples across geographies, based on their physiological class. A principal coordinate analysis (PCoA) was performed on the percent normalized taxonomic profile comprising of 588 genera which were obtained from MG-RAST server (MG-RAST identifier MGP85987 - see Materials and Methods for details) to ensure that the segregation of the metagenomic samples were not dominated by the location of origin. Results of this analysis (Supplementary Figure 1) indicated that the health status of the cows indeed determined the clustering pattern of the bovine milk metagenome. RCM samples were found to cluster closely with CM samples, while H samples formed distinct clusters. The SCM samples were however found to be more closely associated with healthy clusters rather than the diseased groups (CM and RCM). This raised a possibility that SCM may be caused by the triggering of pathogenic mechanisms in some opportunistic pathogen which is otherwise a constituent of the healthy milk microflora, possibly due to changes in the microenvironment. Alternatively, there could be an additional set of pathogens which might determine the severity of mastitis in case of RCM and CM infections.

### Lower abundance of Acinetobacter and Salmonella in milk metagenome could be linked to milder symptoms in SCM

The taxonomic abundance data was used to identify the discriminatory taxonomic groups which were differentially abundant among the microbiota samples in milks collected from cows of varying disease status. Overall, seven taxonomic groups including *Bacillus*, *Catenibacterium*, *Coprobacillus*, *Eubacterium*, *Neisseria*, *Streptococcus* and ‘unclassified genera of bacteria’ were found to be depleted in milk from cows with clinical as well as sub-clinical symptoms (i.e., within the CM, RCM and SCM groups) when compared to healthy cows (Figure 1A). Compared to the healthy samples, the SCM samples were further depleted in the abundance of several taxonomic groups, such as *Acinetobacter*, *Burkholderia*, *Pseudomonas* and *Psychrobacter*. Both CM and RCM metagenome was characterized by a higher abundance of *Klebsiella* w.r.t. H. It was however fascinating to note that the fold-change in the abundance of *Klebsiella* w.r.t. H was higher in RCM (log2FoldChange of 5.613) as compared to CM (log2FoldChange of 3.688), thereby raising the possibility of the involvement of drug resistant *Klebsiella* in RCM infections.

**Figure 1:**
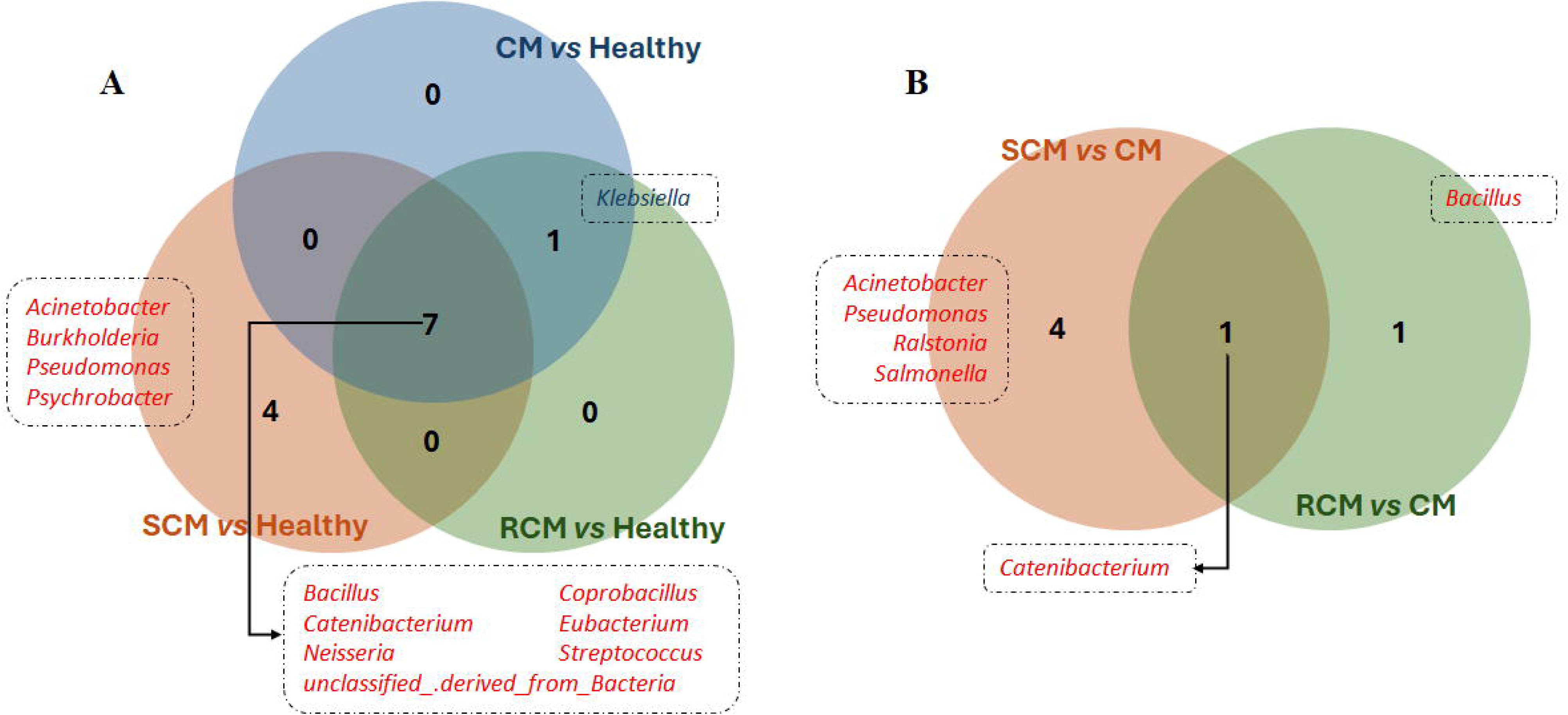
Eular diagrams representing the differentially abundant taxonomic groups in milk microbiota collected from - A. cows having varying disease status/ severity w.r.t. healthy cows (H), and B. cows having recurrent (RCM) and sub-clinical mastitis (SCM) w.r.t. cows suffering from clinical mastitis (CM). While depleted taxonomic groups are indicated in red font, enriched taxonomies are indicated in blue font.

Among the milk metagenomic samples collected from cows suffering from mastitis of varying severities, both SCM and RCM samples were depleted for *Catenibacterium* w.r.t. CM (Figure 1B). In RCM, a lower abundance of *Bacillus* w.r.t. CM was also observed. SCM samples additionally lacked in the abundance of a few bacterial groups like *Acinetobacter*, *Pseudomonas*, *Ralstonia* and *Salmonella* w.r.t. CM. Of these, *Acinetobacter* and *Salmonella* are known to be associated with severe cases of BM (Supplementary Table 2). Hence, their absence/ lower abundance in the milk metagenome could be contributing to the milder sub-clinical symptoms in SCM cases.

### Acinetobacter shares an antagonistic relationship with other bovine pathogens

The taxonomic abundance data was further used to analyse the community interaction patterns among the constituent microbes of H and CM sub-groups (see Materials and Methods). While the CM network comprised of 99 edges representing interactions between 54 genera, the H network was observed to have 74 interactions among 13 genera (Figure 2). The CM network was also observed to be disjoint comprising of two major sub-networks, whereas H network was a single coherent unit with high degree of interaction among the microbes. This close association of microbes within the H network can also be ascertained by the characteristic path length of 1.051 and an average number of neighbours of 11.385. The corresponding values for CM were 2.268 and 4.333 respectively. Nodewise network properties have been provided in Supplementary Table 3. Furthermore, unlike the H network, the CM network reported several negative associations. Most of these negative edges involved *Acinetobacter*, a known bovine pathogen (Supplementary Table 2). The second (major) sub-network in CM network (which was missing in the H network) comprised primarily of bovine pathogens many of which are known causative agents of mastitis (Supplementary Table 2). Among them, *Citrobacter* and *Enterobacter* demonstrated the highest degree and betweenness centralities (Supplementary Table 3). *Neisseria*, *Eubacterium*, *Streptococcus* and *Catenibacterium* were noted to drive the changes in the microbial interactions while analysing the most common sub-network between H and CM networks (Supplementary Figure 2). Further this community shuffling (among the nodes pertaining to the most common sub-network of H and CM networks) was characterized by community splits from ‘control’ – H to ‘case’ – CM (Figure 3). In the three sub-communities thus formed in the CM network, the betweenness centralities of *Neisseria*, *Eubacterium* and *Streptococcus* were noted to increase. Pertinently, among the interactions unique to the CM network, both *Neisseria* and *Eubacterium* were found to interact with *Helicobacter* and *Ralstonia*, both of which harbour pathogenic properties (Figure 2). On the other hand, although *Streptococcus* exhibited a negative interaction with *Acinetobacter*, it had positive interactions with other pathogens like *Ralstonia* and *Granulicatella*. Furthermore, alike *Streptococcus*, *Ralstonia* also exhibited positive interactions with opportunistic pathogens like *Neisseri*a and *Coprobacillus* while being involved in negative interactions with *Acinetobacter*. The above observations highlight the antagonistic relationship between *Acinetobacter* and other bovine pathogens. The probable role of these interactions in the manifestation of bovine mastitis have been elaborated in the Discussion section.

**Figure 2:**
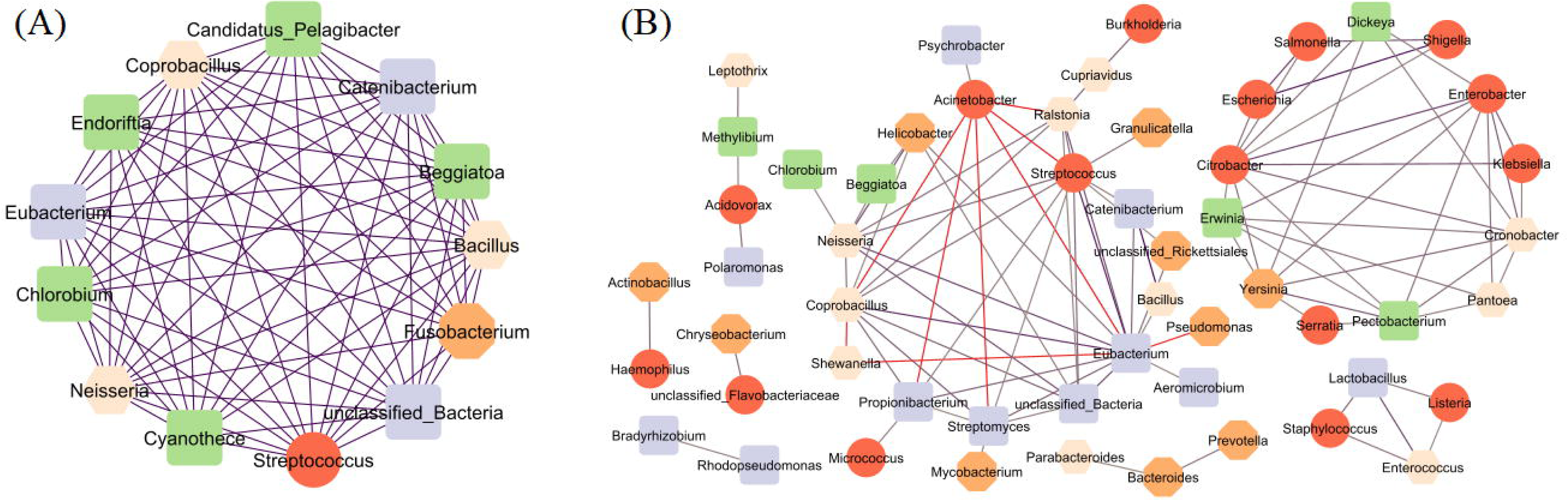
The community interaction network of the microbes derived from milk samples obtained from A. healthy cows (H), and B. cows suffering from clinical mastitis (CM). Nodes representations: red - pathogens known to cause mastitis, orange – pathogens but not known to cause mastitis, pink – opportunistic pathogens, blue – beneficial or commensal microbes, and green – environmental or plant related microbes with or without pathogenic properties. Edge representations: blue (positive interactions) to red (negative interactions) colour gradient representing magnitude of association/ strength of interaction.

**Figure 3:**
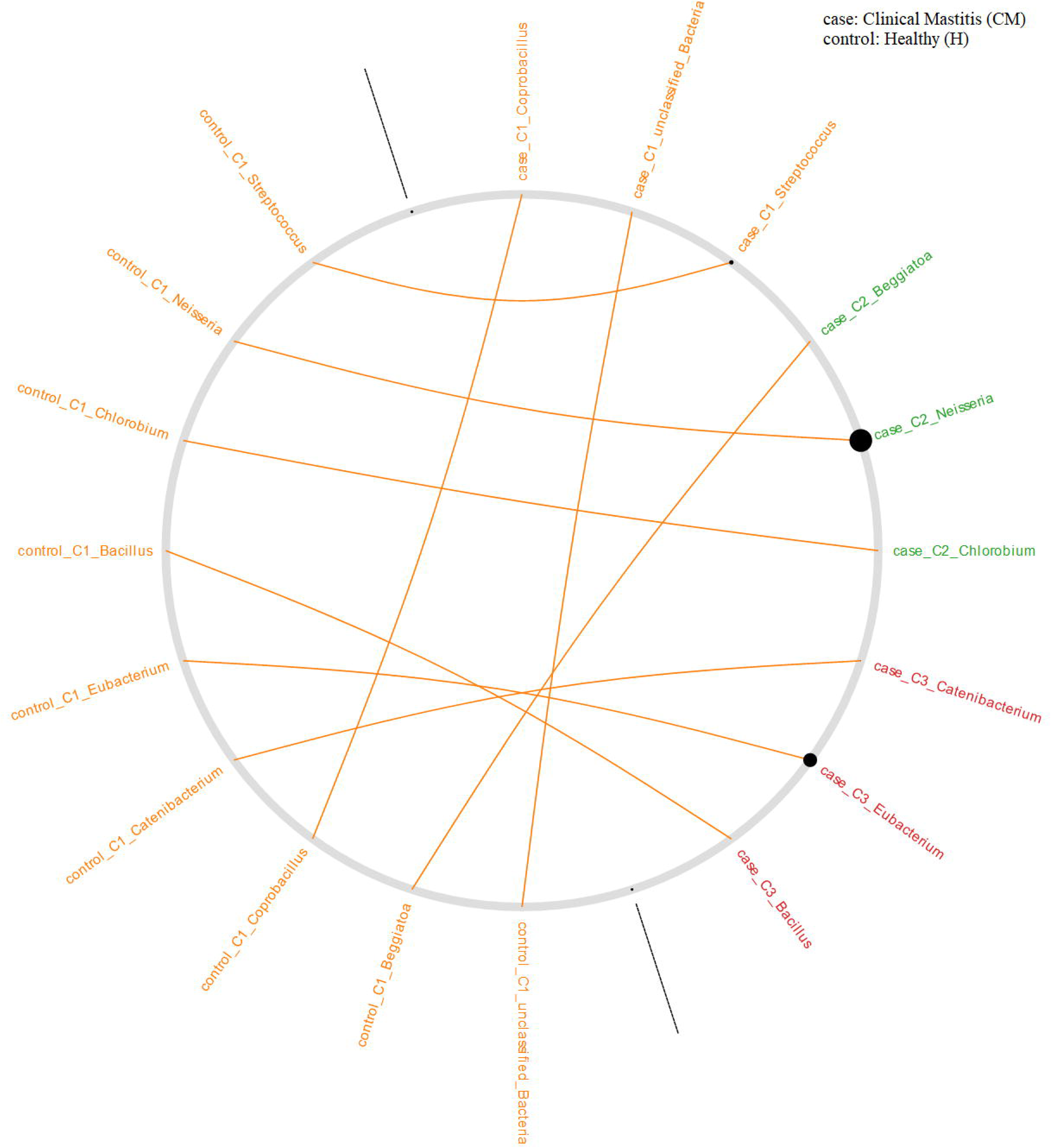
Community shuffle plot depicting the exact changes of the community members (nodes) between ‘control’ – healthy (H) to ‘case’ – clinical mastitis (CM) networks. The plot is displayed as a circle with an axis dividing it vertically into two halves. The left half corresponds to the ’control’ and the right half to the ’case’ sub-network. Node labels in each half corresponds to community affiliation of nodes in the respective networks (indicated by a different colour). Nodes sizes correspond to node betweenness centrality in the corresponding network. Edges connect similar nodes between the two halves (control and case) for ease of visualizing the community shuffling.

### Sensory Protein Load is indicative of sub-clinical symptoms

In line with the analysis performed for identifying discriminatory taxonomic groups from the taxonomic abundance data, the Sensory Protein Load (SPL) profiles were also used to identify distinguishing features among the milk microbiota samples, collected from cows of varying disease status. We could not find any significant variation in terms of SPL among H, CM and RCM sub-groups. However, the SCM sub-group differed significantly in terms of SPL from the other three sub-groups. SPLs of *Achromobacter*, *Dickeya*, *Pectobacterium* and *Raoultella* were identified to be the discriminatory features (Table 1).

**Table 1:** Discriminatory taxonomic groups in terms of sensory protein load (SPL) in milk metagenome of cows with sub-clinical mastitis (SCM) with respect to cows with clinical mastitis (CM), recurrent clinical mastitis (RCM) and healthy (H).

| Taxonomy<br>(SPL-based) | base Mean | Change in SCM w.r.t. CM |  | Change in SCM w.r.t. RCM |  | Change in SCM w.r.t. H |  |
| --- | --- | --- | --- | --- | --- | --- | --- |
|  |  | log2FoldChange | adjusted<br>p-value | log2FoldChange | adjusted<br>p-value | log2FoldChange | adjusted<br>p-value |
| Achromobacter | 137.82 | -23.731 | 6.45E-11 | -24.846 | 3.73E-08 | -23.481 | 7.73E-08 |
| Dickeya | 139.45 | -24.012 | 8.54E-09 | -24.612 | 1.54E-06 | -23.527 | 2.60E-06 |
| Pectobacterium | 150.27 | -24.238 | 1.07E-09 | -24.783 | 3.55E-07 | -22.983 | 1.42E-06 |
| Raoultella | 126.63 | -23.212 | 1.48E-08 | -25.848 | 2.64E-07 | -21.946 | 7.32E-06 |

The Pfam domains associated with each of the sensory proteins contributing to the SPL profile was used to quantify the sensory protein associated Pfam domains (SPPfD) present in the metagenomic samples (see Materials and Methods). The analysis provided 30 unique SPPfDs which were significantly different in abundance between any two of the sample classes (Supplementary Figure 3). Most of the identified SPPfDs were found to be discriminatory between the CM and RCM categories. This included domains like Cache_3, Cache_2, sCache_2, dCache_2, Pkinase, etc. APH domain was exclusively observed to be significantly differential between H vs CM, while IncA domain was significantly differential between H vs CM as well as CM vs RCM. PEGA domain was significantly differential between H vs RCM and RCM vs CM. Domains such as SBP_bac_3, PBP_dimer, Transpeptidase, CHASE and sCache_3_3 was observed to be significantly differential between RCM vs SCM and RCM vs CM. Notably, the taxonomic differences in SCM with respect to CM, RCM or H was not evident from the Pfam analysis. In contrast, the analysis highlighted the differences in functional potential of the RCM metagenome compared to the other three groups.

### Machine learning and model-based prediction

Both the Refseq-based percent normalised profile of taxonomic abundance and the SPL profile (at genus and species levels) were used to train multiple models which were subsequently evaluated upon test set of samples to analyse the efficacy of these models. The models were 100-fold cross validated to prevent any sampling bias during data partitioning. Further, 50 random iterations were carried out to generate these models for each type of profile mentioned above. The median results of these 50-runs have been represented in Supplementary Table 4 for statistical comparison. While accuracy of prediction was highest for models based on SPL profiles cumulated at genus level (92%), species level SPL profile-based models and Refseq based models performed with about 89% accuracy. However, sensitivity and specificity were both observed to be highest for the SPL based models (both at genus and species levels) 0.93 and 1.0 respectively, while the Refseq based models have a sensitivity of 0.87 and specificity of 1.0. The AUC of Refseq based models was 0.9 while for the genus level SPL models it was 0.92, both being higher than the species level SPL models (0.88). Overall, all the ML-models could detect cases of BM with decent accuracies.

## Discussion

Evidence from epidemiological research confirm the nutritional importance of milk and dairy products and even reinforces the possible role of milk in the prevention of several chronic diseases like cardiovascular diseases, some forms of cancer, obesity, and diabetes (Pereira, 2014). Bovine (cow) milk is by far the most consumed variety of milk worldwide accounting for nearly 85% of the global milk production (Claeys et al., 2014). One of the key factors that negatively affects bovine milk yield is mastitis (Ajose et al., 2022; Cheng and Han, 2020). Recent reports suggest that bovine mastitis (BM) is a cause of significant economic loss to the dairy industry with the small and medium-sized farms being most affected (Azooz et al., 2020; Cheng and Han, 2020; Guimarães et al., 2017; Romero et al., 2018). This situation is expected to worsen further as reports suggest that the incidences of BM are likely to increase owing to climate change and global warming (Guzmán-Luna et al., 2022; Vitali et al., 2020). Therefore, timely detection of BM and its clinical sub-types is of immense importance for dairy economics as well as global food security.

The bovine milk microbiota, which closely resembles the cow’s udder microbiota (Taponen et al., 2019) provides a scope for developing periodic screening/ surveillance tools for timely detection of BM. Early detection of sub-clinical mastitis, which is largely asymptomatic, is possibly the biggest challenge. With this aim in mind, a dataset comprising of 42 bovine milk metagenomic samples (Supplementary Table 1) which was collected from five geographical regions (viz., Chittagong, Dhaka, Gazipur, Manikgonj and Sirajgonj) and from cows with varying severities of mastitis (viz., healthy – H; clinical mastitis – CM; sub-clinical mastitis – SCM; and recurrent or chronic mastitis – RCM) was re-analysed in this work. While the geographical origin of the samples did not predominantly affect the cluster patterns, the milk metagenomic samples from healthy cows were found to be more closely associated with the SCM samples and most distinguishing from the CM samples (Supplementary Figure 1). This raised the possibility that the sub-clinical mastitis may be a result of the activation of pathogenic mechanisms in some opportunistic bacteria, which otherwise is a constituent of a healthy milk microflora. Alterations in microenvironment due to the lowered abundance of some of the competing taxonomic groups (Figure 1 and Table 1) might favour these changes. Loss of microbial diversity in SCM by an earlier study (Hoque et al., 2020b) is also indicative of such alterations in microenvironment. Furthermore, the lower abundance/ absence of major mastitis causing pathogens particularly *Acinetobacter* and *Salmonella* in SCM also substantiate the above possibility and indicates that SCM may have a different pathological origin when compared to CM or RCM.

While the SCM and RCM networks could not be derived due to lack of sufficient samples, the H and CM networks were observed to be significantly distinguishing wherein *Neisseria*, *Eubacterium* and *Streptococcus* were observed to drive the changes in community interaction from the healthy to the disease state (Figure 3). We further noted an antagonistic relationship between *Acinetobacter* and other bovine pathogens, particularly *Streptococcus*. It may be anticipated that the manifestation of the symptoms in CM, at least in part, will be driven by the competition between these two causative pathogens and their interaction with other microbes. The majority of the causative agents of BM (which were part of the second sub-network), viz., *Citrobacter*, *Enterobacter*, *Escherichia*, *Klebsiella*, *Salmonella*, *Serratia* and *Shigella* were seen to be very closely associated in the CM network. In this context, it was also fascinating to note the relatively higher abundances (log2foldchange) for all the above-mentioned pathogens (except *Salmonella*) in RCM samples when compared to CM samples (data not shown). However, these observations were not statistically significant – possibly due to lower number of RCM samples. Pertinently, many of them are drug-resistant pathogens (Hoque et al., 2020a) and may possibly explain the recurrence disease phenotype in RCM.

While cases of CM are easier to diagnose, screening/ diagnosis of SCM poses a challenge due to its asymptomatic nature. ML-models to segregate BM affected milk samples from healthy milk samples were therefore built to check if such models can efficiently diagnose all forms of BM including CM, RCM as well as SCM. While the models were created using metagenomic features from CM and H samples only, the independent test set comprised of both RCM and SCM samples along with CM samples. The accuracy of the ML-models (Supplementary Table 4) was indicative of sufficient variations in the metagenomic signatures of the healthy samples from all the three disease categories. However, considering that the sample size used in this study was relatively small, probing of larger datasets is expected to improve the confidence associated with the above predictions.

Overall, our results indicate that the probably pathological mechanisms leading to SCM may be different when compared to CM and RCM. Further, RCM samples may have a slightly higher prevalence of drug-resistant pathogens with respect to CM. Despite these, the metagenomic signatures of all the sub-types of BM are substantially different from healthy milk metagenome. This may be exploited to develop sophisticated ML-models which would allow periodic screening of cows to track the onset of all forms of BM. Given the imminent threat of global warming, this could help in significantly lowering the economic burden of BM on the dairy industry, which in turn would also positively impact the global food security.

## Materials and Methods

### Data Acquisition

The shotgun sequenced bovine milk metagenomic data was obtained from NCBI BioProject PRJNA529353) (Hoque et al., 2020b, 2020a, 2019). The data comprised of 42 bovine milk samples from five different geographical locations in Bangladesh (viz., Chittagong, Dhaka, Gazipur, Manikgonj and Sirajgonj), and four different health status (viz., healthy – H; clinical mastitis – CM; sub-clinical mastitis – SCM; and recurrent or chronic mastitis – RCM). Supplementary Table 1 shows the distribution of the samples across geographies, based on their physiological states. The corresponding taxonomic abundance profiles (at genera level) were also obtained from MG-RAST (MGP85987).

### Construction of Interaction Networks

The taxonomic abundance, in terms of Refseq identifiers (as obtained from MG-RAST), was used to analyse the community interactions for healthy (H) and clinical mastitis (CM) samples. The network of the interacting genera was derived using NetCoMi package in R (Peschel et al., 2021). The SparCC (Sparse Correlations for Compositional data) function within the package was used for estimation of the linear Pearson correlations between the log-transformed components (Friedman and Alm, 2012). The analysis was carried out with alpha set at 0.05, t-test sparsification method, Benjamini-Hochberg correction, no zero filtering, no sample filtering and seed set at 999. While constructing the interaction network for a particular disease status, only the taxonomic groups which were present in at least 50% of the sample in that group were considered. While Cytoscape (Shannon et al., 2003) was used for visualizing the microbial association networks, NetShift (Kuntal et al., 2019) was used to identify the microbes which drove the changes in community interaction between H and CM sub-types.

### Computation of Sensory Protein Load

The sensory protein (SP) related information was obtained from the Sensory Protein Database (SPD) hosted at https://osf.io/gr6h5/ (Bhar et al., 2022, 2021). Sequence alignment, using BLAST (Camacho et al., 2009), of the metagenomic reads were performed against the SPD to obtain quantitative profiles of SP content in each of the metagenomic samples. The best hits qualifying a minimum e-value cutoff of 1e^-5^ were considered. Subsequently, the Sensory Protein Load (SPL) (Bhar et al., 2022) for each of the species in each of the metagenomic samples were computed. The cumulative aligned length (covered base length) of the hits for a species (in each metagenomic sample) indicated the total length of the potential SP coding regions in the metagenome for that species (in that sample). The sensory protein load (SPL) was hence calculated using the following formula:

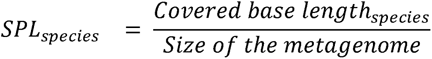

where, the size of the metagenome is the cumulative length of the reads constituting the metagenome (in megabase units). SPL for genus level was computed by cumulating the SPLs at species level.

### Analysis of Discriminatory Taxonomic Groups

The non-normalized taxonomic abundance data at genera level, in terms of Refseq identifiers (as obtained from MG-RAST), was used for discriminatory analysis using the DESeq2 package in R (Love et al., 2014). The healthy samples (H) were used as the reference level while contrast was drawn between, RCM vs CM and SCM vs CM. Similarly, the SPL signatures (at genera level) was also subjected to DESeq2 analysis to identify those microbes which encoded for differential abundance of SPs on their genome, between the different sample groups (viz., H, CM, RCM and SCM) while keeping healthy samples (H) as the reference. Default parameters were used for DESeq2 except for the sfType being “poscount”, test performed being “Wald”, alpha kept at 0.05, and pAdjustMethod being “fdr”. Taxonomic groups qualifying the criteria of baseMean ≥ 100, log2FoldChange ≥ ±2 with an adjusted p-value < 0.05 were considered as significantly discriminatory taxonomic groups.

### Pfam Domain Analysis

In alignment with our previous study, the sensory protein associated Pfam domains (SPPfDs) for each metagenomic sample were quantified (Bhar et al., 2022). The obtained SPPfD values were further normalized by the metagenome size and SPPfDs found to be present in at least 50% of the samples of any one of the heath classes/ disease sub-types were considered for further analysis. For each of these SPPfDs, pairwise Wilcoxon test between the sample classes were performed and the domains which were found to be significantly differential (p-value ≤ 0.05) in any of these pairwise comparisons were considered significant. The significantly differential SPPfDs for all the pairwise class comparisons were plotted with help of the packages ggplot2 and ggpubr (Kassambara, 2023; Wickham et al., 2019).

### Construction of Early Risk Assessment Models

The taxonomic abundance profile obtained from MG-RAST (post percent normalization) was used to check if a ML-based classifier models for early risk assessment of bovine mastitis can be constructed. Models were trained on 80% of the samples (called “training set”) belonging to healthy and CM class only, while the rest 20% of H and CM, as well as all the SCM and RCM samples (known as “test set”) were kept exclusively for testing. During training of the model, the “training set” was further randomly split into an internal training (80%) and an internal testing (20%) set. Given the relatively lower number of samples in the H category compared to the CM category, synthetic minority oversampling technique (SMOTE) was applied to remove any potential bias (Chawla et al., 2002). An initial feature selection through the recursive feature elimination (Kuhn, 2008) was employed and the Random Forest algorithm (Liaw and Wiener, 2002) was used for prediction. The model was subjected to 100-fold cross validation. Further, the entire workflow was repeated for 50 random iterations to rule out any chance of sampling bias arising from partitioning of the data. Similarly, models were built using the SPL profile (both at genus and species levels) as well. Statistical comparisons were performed using pROC package (Robin et al., 2011) in R.

## AUTHOR CONTRIBUTIONS

TB designed the study. SB performed the analysis. SB and TB interpreted the data and wrote the manuscript.

