## Supplementary figures and images for "Evidence from metagenomic study indicate that subclinical mastitis may have a different pathological origin than clinical mastitis"

### Supplementary Figure 1

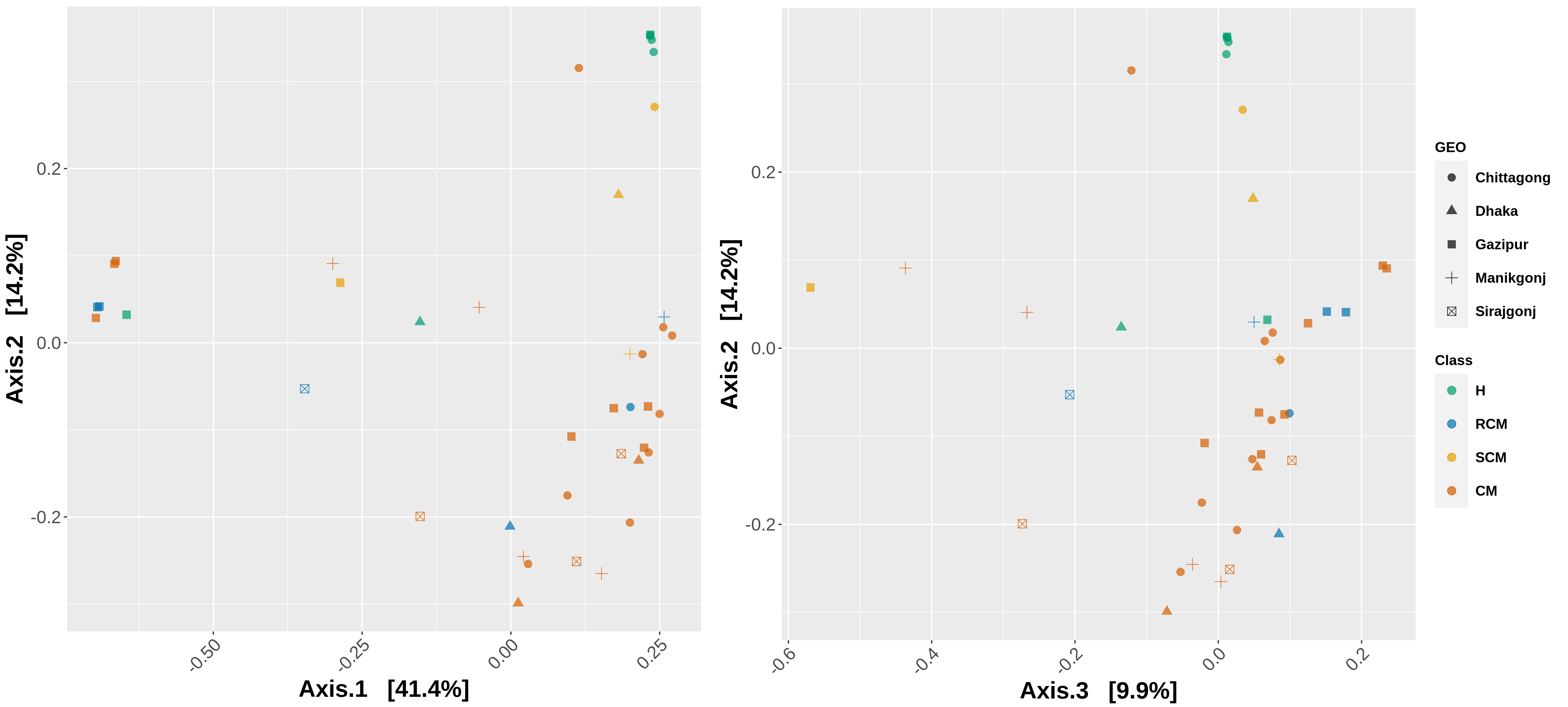

### Supplementary Figure 2

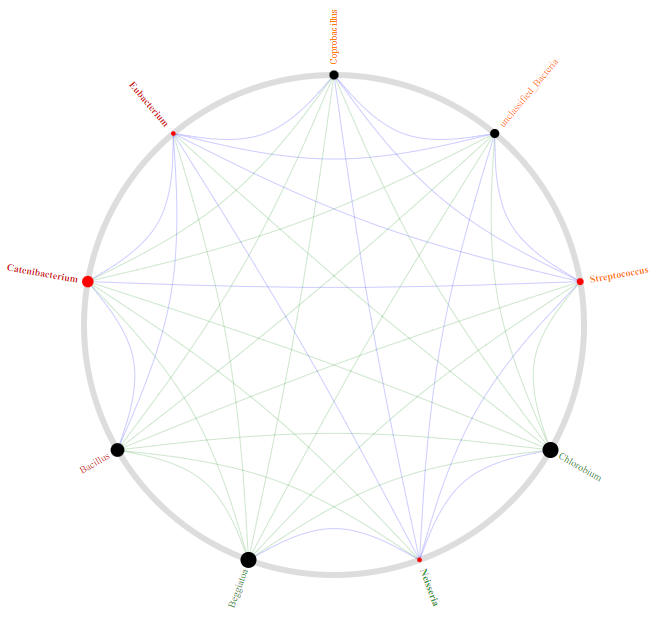

### Supplementary Figure 3

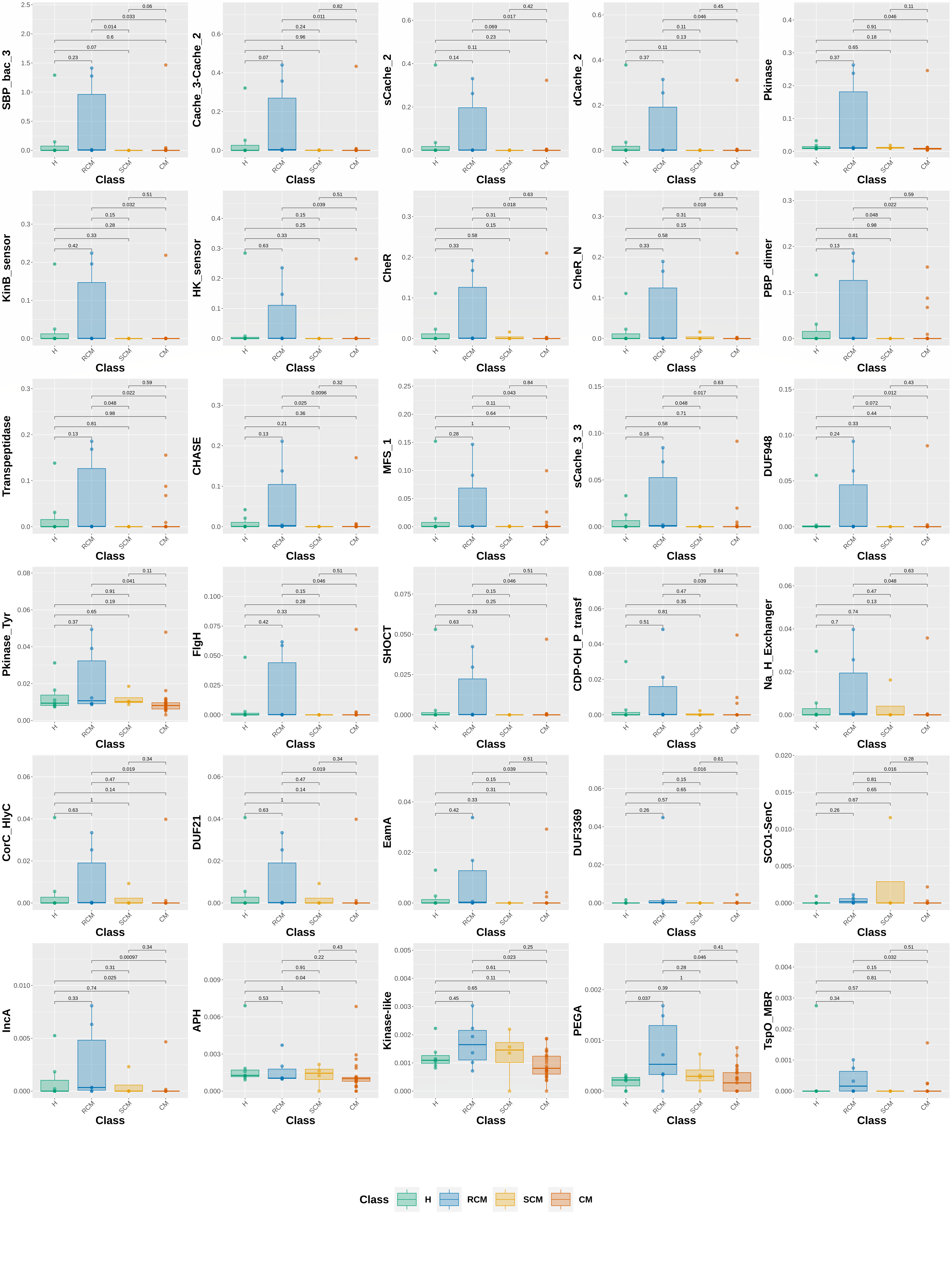
